# Impaired dendritic spine development in a zebrafish model of Fragile X

**DOI:** 10.1101/2022.02.15.480580

**Authors:** Elisabeth C. DeMarco, George Stoner, Estuardo Robles

**Affiliations:** Department of Biological Sciences and Purdue Institute for Integrative Neuroscience, Purdue University, West Lafayette, IN, USA

**Keywords:** Neuronal development, optic tectum, synaptogenesis

## Abstract

Dendritic spines are the principal site of excitatory synapse formation in the human brain. Impaired formation of spines during development has been observed in several autism spectrum disorders (ASDs), including Fragile X syndrome. Fragile X is caused by transcriptional silencing of the *Fmr1* gene encoding the RNA-binding protein FMRP (Fragile X mental retardation protein). While spine development has been well characterized in the mammalian CNS, spines are not unique to mammals. Pyramidal neurons (PyrNs) of the zebrafish optic tectum form an apical dendrite containing a dense array of dendritic spines. We employed a genetic labeling system to monitor PyrN dendritic spine development in larval zebrafish. Our findings identify a developmental window when PyrN dendrite growth is concurrent with spine formation. Throughout this period, motile, transient filopodia gradually transform into stable spines containing postsynaptic specializations. *fmr1* mutant zebrafish larvae exhibit pronounced defects in both PyrN dendrite growth and the formation of morphologically mature spines. Live imaging of PyrN dendrites suggests these defects are caused by an inability to stabilize nascent contacts. These findings indicate spine stabilization is essential for PyrN dendritic arborization and establish zebrafish larvae as a model system to study spine development *in vivo*.

## Introduction

Dendritic spines are the principal site of excitatory synapse formation in the human brain (Nimchinsky et al., 2002; Yuste, 2013). The developmental processes leading to spine formation have been well characterized in the mammalian brain. Following the establishment of neuronal polarity, dendrites grow and branch to form an arbor defining the neuron’s synaptic input field (Ledda and Paratcha, 2017). Following this phase, the arbor is remodeled via activity-dependent addition or removal of branches. Spine development is then initiated by formation of filopodia from the dendritic shaft (Ziv and Smith, 1996). Filopodia are motile, antenna-like protrusions that sample the environment for extracellular cues and potential synaptic partners. Contact with a synaptic partner leads to local recruitment of postsynaptic density (PSD) scaffolding proteins such as PSD95, a membrane-associated guanylate kinase (MAGUK) with a role in localizing glutamate receptors to PSDs (Chen et al., 2015). Formation of a synapse stabilizes the filopodium and initiates its transformation into a mature spine with an enlarged head containing a PSD (Harris, 2020).

Changes in the morphology and/or density of dendritic spines is a common neuropathology associated with several autism spectrum disorders (ASDs). Histological examination of post-mortem tissue from Fragile X patients revealed a dramatic change in spine morphology typified by a prevalence of “immature, long, tortuous spines” (Hinton et al., 1991). Consistent with these findings, immature spine morphologies have been described in several brain areas of *Fmr1* knockout mice, including cortex (Comery et al., 1997; Cruz-Martín et al., 2010; Nimchinsky et al., 2001; Pan et al., 2010), hippocampus (Grossman et al., 2006), cerebellum (Koekkoek et al., 2005), and amygdala (Qin et al., 2011). Although several studies in cortex reported changes in spine density, more recent studies have reported no significant spine density changes in *fmr1* knockout mice (Cruz-Martín et al., 2010; Harlow et al., 2010; Pan et al., 2010; reviewed by He and Portera-Cailliau, 2013). While these discrepancies may be due to differences in developmental timepoint examined or imaging method, cell types likely differ in their sensitivity to loss of the *fmr1* gene (He and Portera-Cailliau, 2013). For example, cerebellar Purkinje cells in *fmr1* knockout mice exhibit normal dendritic arbor complexity and spine density while forming spines with immature spine morphologies (Koekkoek et al., 2005). Together, these studies point to impaired spine maturation as the most consistent defect in *fmr1* knockout mice.

Dendritic spines are not unique to the mammalian brain – dendritic spines are also found in coldblooded vertebrates and invertebrates. Remarkably, structures resembling dendritic spines are common among neurons in the nervous system of planarians, the simplest living animals with a bilateral body plan (Sarnat and Netsky, 1985). Although rare in insects, honeybee Kenyon cells in the mushroom body have dendrites densely decorated with spine-like protrusions (Groh and Rössler, 2020). Kenyon cell spine densities remain constant during development, yet exhibit a shift towards shorter and thicker morphologies (Farris et al., 2001). In the fruit fly, lobula plate tangential cell dendrites form spines resembling vertebrate spines in shape, size, and density (Leiss et al., 2009). Of note, these appear to be a site of cholinergic neurotransmission, whereas the vast majority of vertebrate spines are sites of glutamatergic neurotransmission. In cold blooded vertebrates, dendritic spines have been described in *Xenopus* olfactory bulb interneurons (Zhang et al., 2016) and in tectal interneurons of the jewel fish (Coss and Globus, 1979). However, these systems lack genetic methods to consistently label a stereotyped population of spiny neurons. In the larval zebrafish, most genetically-identified tectal neuron types lack dendritic spines (Del Bene et al., 2010; Helmbrecht et al., 2018; Kunst et al., 2019; Robles et al., 2011; Scott and Baier, 2009). This is surprising considering 1) PSD structure and protein composition is well conserved across vertebrate species (Bayés et al., 2017) and 2) dendritic spines with PSDs have been observed in the tectum of adult zebrafish, goldfish, and perch (Bayés et al., 2017; Laufer and Vanegas, 1974; Meek, 1981).

One neuron type in the adult teleost tectum known to form dendritic spines is the Type I/pyramidal neuron (PyrN) (Folgueira et al., 2020; Ito and Kishida, 2004; Meek, 1990; Vanegas et al., 1974; Xue et al., 2003). We previously described an *id2b:gal4* transgenic that labels PyrNs in the larval zebrafish tectum (DeMarco et al., 2019). A subsequent study used sparse genetic labeling to demonstrate that PyrNs, despite forming small synaptic territories, are densely innervated with as many as 100 postsynaptic specializations (Demarco et al., 2021). One PyrN structural feature that facilitates this high input density is the presence of dendritic spines along its apical dendrite (Demarco et al., 2021; Folgueira et al., 2020; Laufer and Vanegas, 1974). These serve to increase dendritic surface area and biochemically isolate individual PSDs. The ability to reliably label a genetically-specified spiny neuron type will allow us to leverage the strengths of the zebrafish model system to examine mechanisms of spine formation. We set out to characterize the process of spine maturation in the larval zebrafish tectum with the goals of 1) identifying an appropriate developmental window to study spine development, 2) monitoring protrusion dynamics during the filopodia-to-spine transition, 3) defining the relationship between PSD95 accumulation and spine development, and 4) determining if PyrN spine maturation is disrupted in *fmr1* mutant larvae.

Our findings identify a developmental window in early larval development from 4-11 days when PyrN dendrite remodeling (branch addition/elimination) is concurrent with spine formation. Throughout this period, motile, transient filopodia are gradually replaced by short, stable spines. During this transition, the postsynaptic protein PSD95 localizes to most spines, yet its accumulation is a weak predictor of spine stability. Structural imaging of PyrN apical dendrites in *fmr1* mutants revealed smaller dendrite arbors with reduced spine densities. Morphologically, these spines had thinner heads compared to those in wild-type larvae. Time-lapse imaging confirmed these spines as less stable and exhibit high rates of turnover. These spine defects are consistent with previous findings in mammals and establish the *id2b:gal4* transgenic as a valuable tool to study spine development in a genetically tractable and optically transparent vertebrate.

## Results

### Tectal PyrNs contain spiny apical dendrites

*In vivo* examination of PyrN development was performed using mosaic genetic labeling by injection of *Tg*(*id2b:gal4,uas-e1b:ntr-mcherry)* transgenic embryos with DNA encoding a membrane targeted EGFP under control of a UAS enhancer region (*uas:egfp-caax*). Injection of 150-200 two-to four-cell stage embryos typically yielded several larvae with strong, sparse labeling of isolated neurons in tectum (Figure 1A-B). The *id2b:gal4* transgenic labels three tectal neuron types: PyrNs (73.5%), torus longitudinalis projecting neurons (11.8%), and tegmentum projecting neurons (14.7%) (DeMarco et al., 2019). In our initial experiments, these three neuron types could be consistently distinguished morphologically by 4 days post-fertilization (dpf). PyrNs were primarily identified by their dendritic stratification pattern, with three arbors formed in different synaptic layers of the tectal neuropil (Figure 1C). The apical dendrite is formed in the superficial-most stratum marginalis (SM) layer and receives excitatory input from the torus longitudinalis (TL), a second order visual area in fish (Figure 1C) (Folgueira et al., 2020; Northmore, 2017). The medial dendrite is formed in the stratum fibrosum grisealis (SFGS) layer and has been shown to receive direct inputs from retinal axons (Laufer and Vanegas, 1974). The basal dendrite situated in the stratum grisealis centralis (SGC) is a mixed neurite arbor containing both axonal and dendritic branches (Demarco et al., 2021). In every PyrN imaged, the developing neuron resembled its mature, tristratified form at 4 days of development (n=9 PyrNs from 8 larvae). High resolution imaging of PyrNs labeled by *id2b:gal4* transgene confirmed numerous spine-like protrusions on the apical dendrite at 8 dpf (Figure 1D). Structurally, these ranged from thin, filopodia-like protrusions to ones with mushroom-shaped heads (Figure 1E). Additionally, we observed branched spines with two distinct heads (Figure 1E). Spines with mushroom or branched heads typically exhibited lengths in the range of 0.5 to 2μm, whereas thin filopodia-like protrusions could be as long as 5μm. In general, the shapes and dimensions of larval PyrN spines were similar to those in mammals (Sheng and Kim, 2011) and adult zebrafish (Bayés et al., 2017).

**Figure 1.**
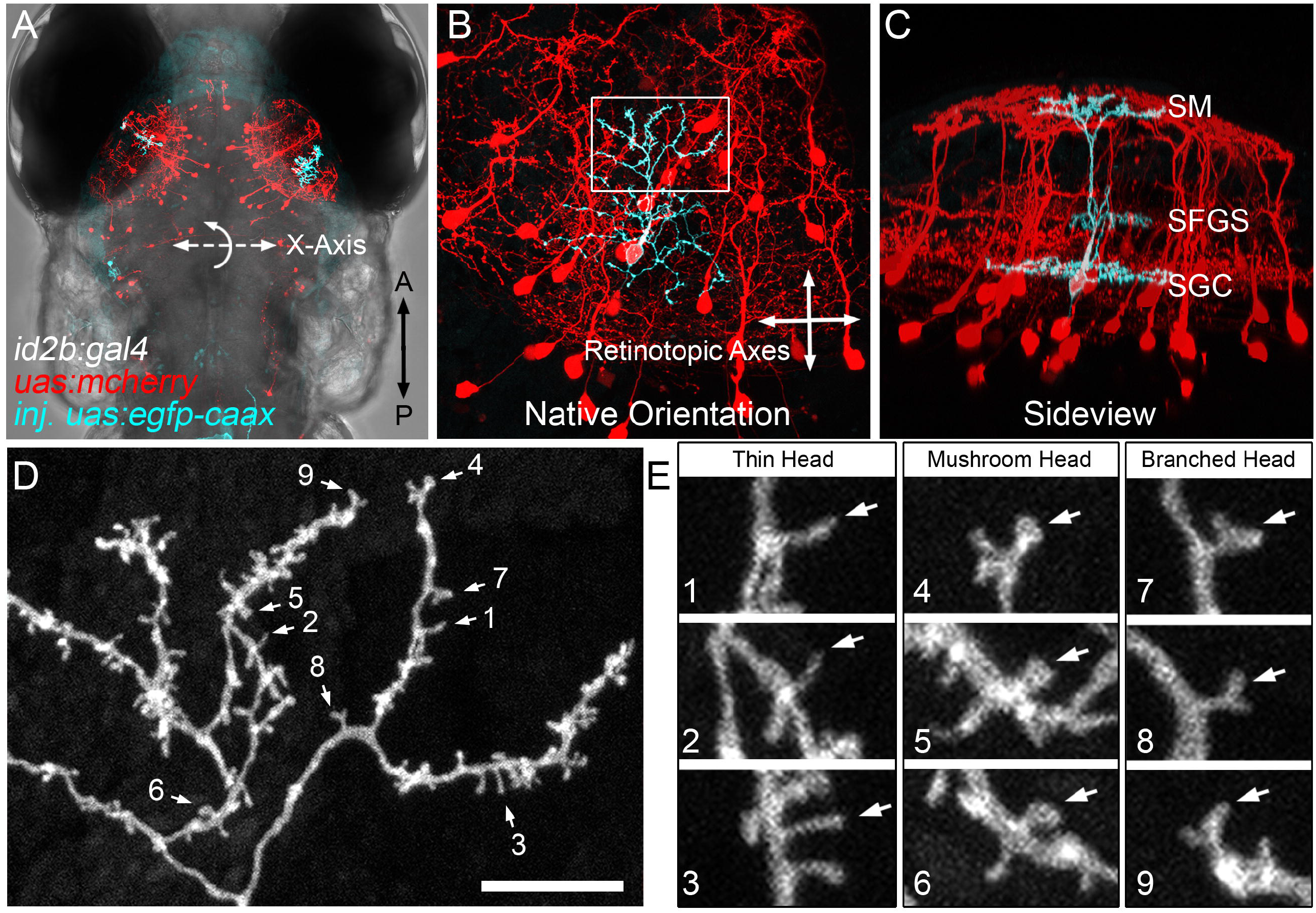
Dendritic spines on PyrN apical dendrites. **A**. Dorsal view, whole-brain confocal image volume of an 8 dpf double transgenic *Tg(id2b:gal4,uas-e1b:ntr-mCherry)* larva injected at the embryo stage with a *uas:egfp-caax* plasmid to generate sparse labeling. Note single EGFP-labeled PyrN in each tectal lobe. **B**. Higher magnification maximum projection of neuron labeled in right tectal lobe of larva in A. Projection is shown from dorsal view, 0º rotation. Note that this view was used for subsequent measurements of dendrite area along the retinotopic axes. **C**. Maximum projection of same neuron rotated -50º about the X-axis, to yield an orientation parallel to the tectal layers. Note clearly stratified neurite morphology with arbors in SM, SFGS, and SGC layers of tectal neuropil. **D**. High magnification view of a subvolume of the SM-targeted dendrite of PyrN in B-C as indicated by box in B. Note the branched arbor decorated with multiple short protrusions. Arrows indicate spines selected for higher magnification views. **E**. 3x magnified views of 9 dendritic spines indicated by arrows in D. Note the presence of different types of spine heads: thin, mushroom, and branched. Scale bar: 200μm in A, 50μm in B-C, 10μm in D, 5μm in E.

### PyrN dendrite stratification precedes dendritic arbor stabilization

To determine the order in which characteristic morphological features of PyrNs are established, we conducted multi-day imaging of single PyrNs labeled with EGFP-caax. Two hallmark structural features of PyrNs are 1) a tristratified dendrite and 2) an apical dendritic arbor containing a dense constellation of spines. We reasoned these two processes should be concurrent if spine formation serves an instructive role in guiding dendrite stratification. In three larvae, we were able to perform longitudinal, multi-day imaging of the same PyrN at 4, 6, 8 and 11 dpf. Confocal image volumes acquired at 4dpf revealed PyrNs with highly branched dendritic arbors (Fig 2A). Between 4 and 11 dpf, the arbor underwent structural rearrangements and appeared to have fewer fine branches (Fig 2B). Sideview rotations of these image volumes enabled visualization of the three characteristic PyrN dendrite stratifications (Fig 2C-D). In every neuron examined, the stratification pattern remained constant from 4 to 11 dpf. During the same period, the vertically oriented primary dendrite branch increased in length due to increasing thickness of the tectal neuropil (Fig 2D). Although the stratification pattern remained largely unchanged, arbors did undergo structural rearrangements along the retinotopic axes (orthogonal to stratification layers). Most notable in the PyrN in Figure 2 was an increase in arbor size between 4 and 6 dpf (Fig 2E-F). Examining the apical dendrite in more detail, it had a highly filopodial morphology at 4 dpf (Fig 2E), whereas protrusions with spine-like shapes predominated at later timepoints (>6 dpf; Fig 2F-H). This confirms that PyrN stratification pattern is established early and remains stable throughout the period of larval development examined (4-11 dpf). PyrN dendrite stratification may be genetically specified and independent of activity dependent refinements controlling arborization along the retinotopic axes. This would be consistent with previous findings that synaptic layering in the tectum is “hardwired” and unaffected by blockade of neuronal activity or neurotransmitter release (Nevin et al., 2008).

**Figure 2.**
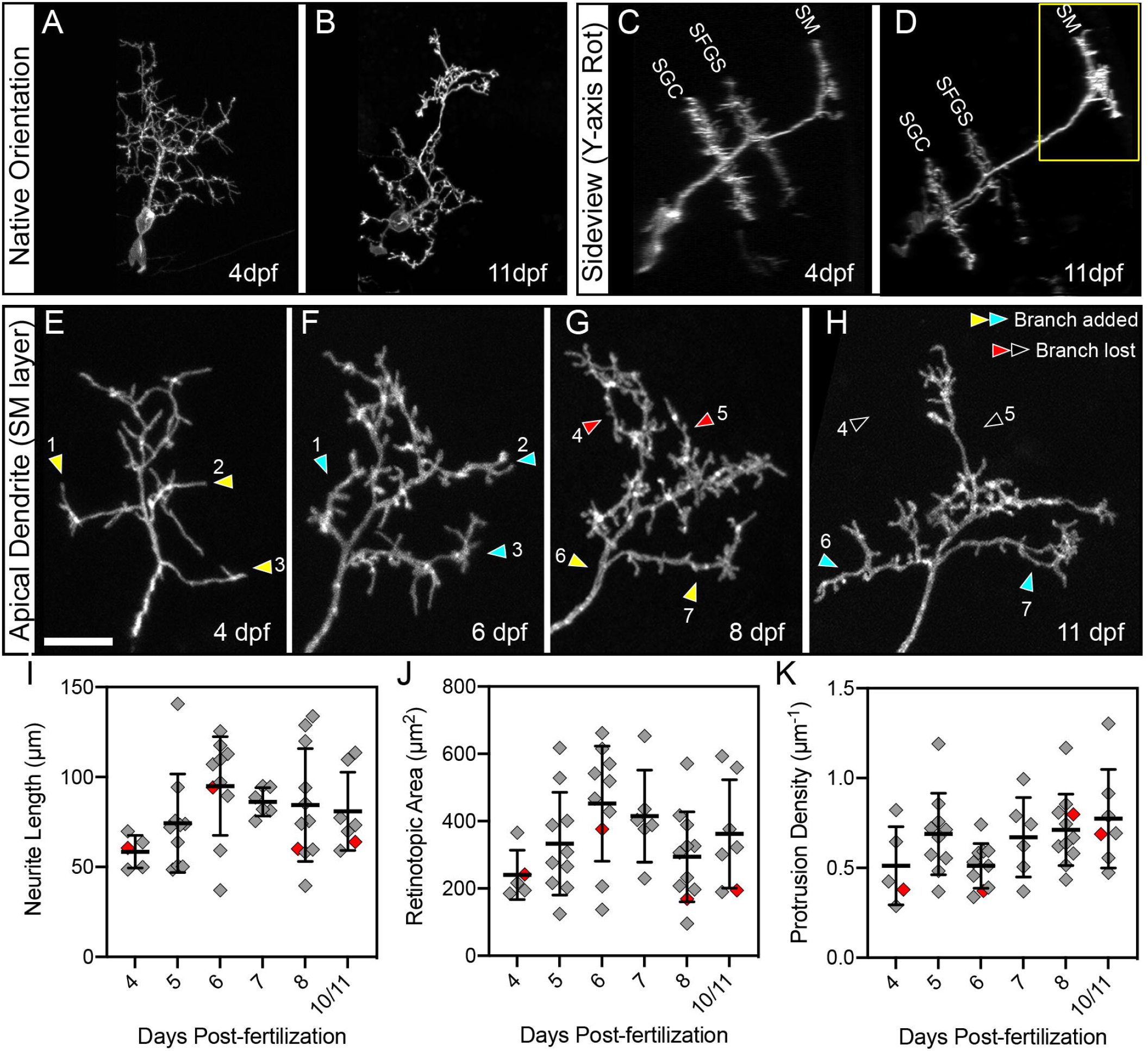
Multi-day imaging of dendrite development and spine formation. **A-B**. Native orientation (dorsal view) maximum projection images of the same PyrN at 4 dpf and 11 dpf. **C-D**. Rotated sideview projections of neuron in A-B. Three distinct dendritic stratifications are visible at each timepoint. Also note that the total length of the main dendritic trunk increases, most likely due to thickening of tectal neuropil during this developmental period. **E-H**. Higher magnification views of the SM-targeted apical dendrite of PyrN in A-D (region indicated by yellow box in D). Sites where new branches are added are indicated by yellow arrowheads (before branch addition) and cyan arrowheads (after branch addition). Sites where branches were retracted are indicated by red arrowheads (before branch retraction) and black arrowheads (after branch retraction). Note that branches are added and retracted throughout this time window. Also note that from 4 to 11 dpf there is a gradual loss of long, filopodial protrusions and an increase in short, spine-like protrusions. **I**. Neurite length measurements of PyrN SM dendrites between 4 and 10/11 dpf. Note that due to reduced survival at 10 and 11 dpf timepoints these groups were combined into one. **J**. Neurite length measurements of PyrN SM dendrites between 4 and 10/11 dpf. **K**. Spine density measurements of PyrN SM dendrites between 4 and 10/11 dpf. Number of larvae analyzed for each timepoint: 5, 10, 10, 6, 11, and 7. All graphs mean ± S.D. Statistical test: one-way ANOVA with Tukey’s multiple comparisons posthoc test, p values for all pairwise comparisons in I, J, and K were >0.05. Scale bar: 20μm in A-D, 5 μm in E-H.

### PyrN apical dendrite arborization is concurrent with the filopodia-spine transition

Synaptotropic models of neurite growth propose that synapse formation guides dendrite arborization (Cline and Haas, 2008; Niell, 2006). As spines are likely the main site of synapse formation on PyrN dendrites, this would require concurrent dendrite growth and spine formation. To determine the time-course of apical arbor stabilization relative to the filopodia-spine transition, we examined image volumes containing the apical dendrite (boxed region in Figure 2D) over multiple days of development. This allowed us to monitor dendritic branches gained and lost over time (see Figure 1b). Figure 2E-H shows the structural development of a single PyrN apical dendrite at 4, 6, 8 and 11 dpf. At 4 dpf, this PyrN has a compact apical dendrite containing several branches and many filopodial protrusions (Figure 2E). Between 4 and 6 dpf, this arbor undergoes an increase in size driven by the addition of three new branches (yellow-to-cyan arrowheads in Figure 2E-F). By 6dpf, this dendrite has a mix of filopodia-like protrusions and short, spine-like protrusions. From 6 to 8 dpf, no branches are gained or lost and the total area is similar (Figure 2F-G), however the majority of protrusions are now short and spine-like. Between 8 and 11 dpf, the arbor gains two branches (yellow-to-cyan arrowheads in Figure 2G-H) and loses two branches (red-to-open arrowheads in Figure 2G-H). Throughout these changes in shape, the dendrite maintained a similar density of spine-like protrusions (Figure 2G-H). This is reminiscent of previous findings in aspiny tectal neurons, where dendrite length and synapse number increase between 3 and 7 dpf but remain constant from 7 to 10 dpf (Niell et al., 2004).

To quantitatively assess PyrN apical dendrite growth during the filopodia-spine transition, we analyzed three morphological parameters between 4 and 11 dpf: total neurite length, retinotopic area, and protrusion density. 3D neurite lengths were calculated by generating skeletonized tracings of the dendrite. Retinotopic area was calculated using a convex polygon connecting the outermost branch tips of each arbor along the retinotopic axes (see Figure 1b). Protrusion density was calculated by manually annotating protrusion locations on maximum projections of dendrite image volumes. Neurite length gradually increased between 4 and 6 dpf, followed by a slight decrease from 6-10 dpf (Figure 2I). A similar pattern was observed when examining the retinotopic area of PyrN apical dendrites area from 4 to 11dpf. However, these trends were not statistically significant (Figure 2I-J; one-way ANOVA with Tukeys multiple comparisons test post hoc). Although their morphology appeared to change over time, dendritic protrusions density remained relatively constant between 5 and 11 dpf (5 vs 6, 7, 8, or 10/11 dpf, all p values >0.418; Figure 2K). This suggests apical dendrites are simultaneously sampling a large number of potential presynaptic partners during early stages of arborization.

To quantify protrusion maturity during this developmental period, we measured two morphological metrics: protrusion length and head width. To minimize the possibility of underrepresenting thinner or dimmer protrusions, we performed these analyses on images generated using a permissive threshold mask so all spines were filled with equal pixel intensities (Figure 3A-D). These analyses included isolated spines located anywhere on the dendritic arbor. In total, length measurements were obtained from 1,043 protrusions (>120 at each timepoint, >5 neurons at each timepoint). From 4 to 11 dpf, the average length of protrusions gradually decreased (Figure 3I), driven by a reduction in the number of protrusions >2 μm. To examine spine head width, we acquired image volumes from 5 and 8dpf PyrNs at increased resolution (pixel sizes between 0.03 and 0.08 μm) by using a higher numerical aperture objective and increasing image size. 5 dpf was chosen as the early timepoint because these PyrN dendrites exhibit a high density of filopodia-like protrusions (Figure 3A, C, and I). 8 dpf was chosen as the late timepoint because the majority of protrusions at this stage were spine-like and <2μm in length (Figure 3B, D, and I). Head widths were measured by obtaining intensity profiles from regions spanning the distal 0.5 μm of each protrusion (Figure 3E-H). From these traces, we analyzed the width of each curve at half-maximum to generate width measurements for >50 spines at each age. Although this approach may slightly underestimate the width, the values obtained were similar to the spine head widths observed in the adult zebrafish tectum (Bayés et al., 2017). Despite a range of widths observed at both time-points, we did detect a significant increase in protrusion head width from 5 to 8 dpf (p<0.001, unpaired T test), reflecting an increase in spine-like morphologies. Concurrent dendrite arborization and filopodia-spine transition is consistent with a model in which spine formation guides dendritic growth and branching.

**Figure 3.**
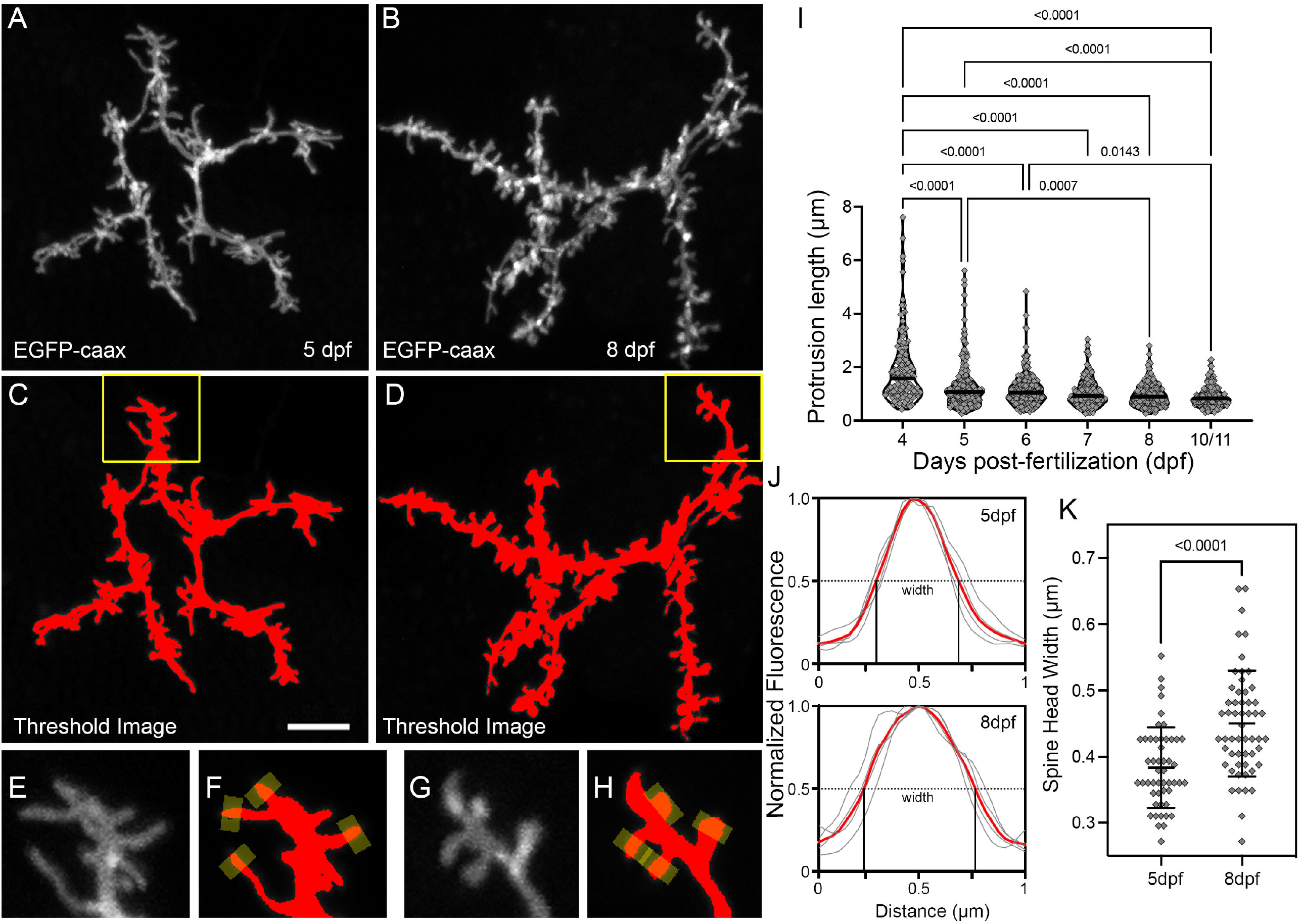
Morphological changes associated with the filopodia-spine transition. **A-B**. Maximum projections of apical dendrite subvolumes acquired from a 5 dpf PyrN (A) and an 8 dpf PyrN (B). **C-D**. Examples of images in A-B with a low threshold mask applied. **E-H**. Magnified views of dendrite subregions indicated by yellow boxes in C and D. Grayscale and thresholded images are presented for each. Yellow rectangles at spine tips indicate 0.5 μm wide line selections used to generate fluorescence intensity plots across each spine head. **I**. Quantification of protrusion lengths for PyrN apical dendrites at 4, 5, 6, 7, 8, and 10/11 dpf. Note gradual decrease in the incidence of protrusions with lengths greater than 2 μm. Median for each group is indicated by horizontal line. Number of spines and neurons analyzed at each timepoint: 123 (6), 225 (7), 214 (7), 137 (5), 187 (5), and 157 (4). One-way ANOVA with Tukey’s multiple comparisons posthoc test. All significant differences (p<0.05) are shown on graph. **J**. Spine head fluorescence intensity plots for measuring spine head widths. Thin gray traces are for the 4 spines indicated in F and H. Width was calculated by measuring the width at half-max for each trace. Mean of these widths for each set is indicated by red trace. **K**. Comparison of spine head widths at 5 and 8 dpf. Data presented as mean ± S.D. Unpaired t-test. Scale bar: 5μm in A-D, 2 μm in E-H.

### Changes in spine morphology correlate with increased stability

Our first indication that dendritic protrusions become more stable during larval development came from timelapse recordings of neurons used for multi-day structural imaging (Figure 2). At 4 dpf, arbors contained motile filopodia dynamically extending and retracting, whereas at 8 dpf most protrusions were spine-like and present for several hours (Supplemental Movie 1). However, even these stable spines present for the entire 4hr recording constantly underwent small changes in shape and length (typically ≤1μm). To qualitatively demonstrate the difference in number of motile protrusions, we generated color-coded temporal projections from dendritic timelapse recordings (Figure 4A-D). These images were made by assigning a unique color to each timepoint and combining these single-color images into a maximum projection image. In the resulting image, motile and short-lived protrusions are labeled with a single color, whereas stable regions appear white (composite of all colors in lookup table). While 4dpf dendrites contained many motile and short-lived protrusions, whereas 8dpf dendrites contained very few (arrowheads in Figure 4B, D). To quantitatively assess the stability of dendritic spines, we directly measured protrusion lifetimes from 4hr timelapse recordings collected at 4, 5, 6, 7, 8, and 10/11 dpf (n=5 neurons and >100 protrusions per timepoint; Figure 4E). At 4 and 5 dpf, stages when PyrN apical dendrites have many filopodia-like protrusions, very few protrusions had lifetimes ≥230min (5.5% and 12.1%, respectively; n=73 and 91; Figure 4E). Conversely, at 7, 8, and 10/11dpf close to half the protrusions had lifetimes ≥230min (40.1%, 48%, and 55.5%, respectively; n=83, 123, and 110; Figure 4E). 6dpf represented an intermediate stage where 34.3% of protrusions were stable. Statistical testing confirmed a significant difference between early (4-5 dpf) and late timepoints (7, 8, and 10/11dpf; p<0.0001 for all pairwise comparisons between early and late, one-way ANOVA with Tukey’s multiple comparisons test). This pattern is the opposite of what we observed for protrusion lengths, which incrementally decreased during this period (Figure 3I). Therefore, the shift from filopodial to spine-like morphologies coincides with an increase in protrusion stability, likely due to stable synaptic contact.

**Figure 4.**
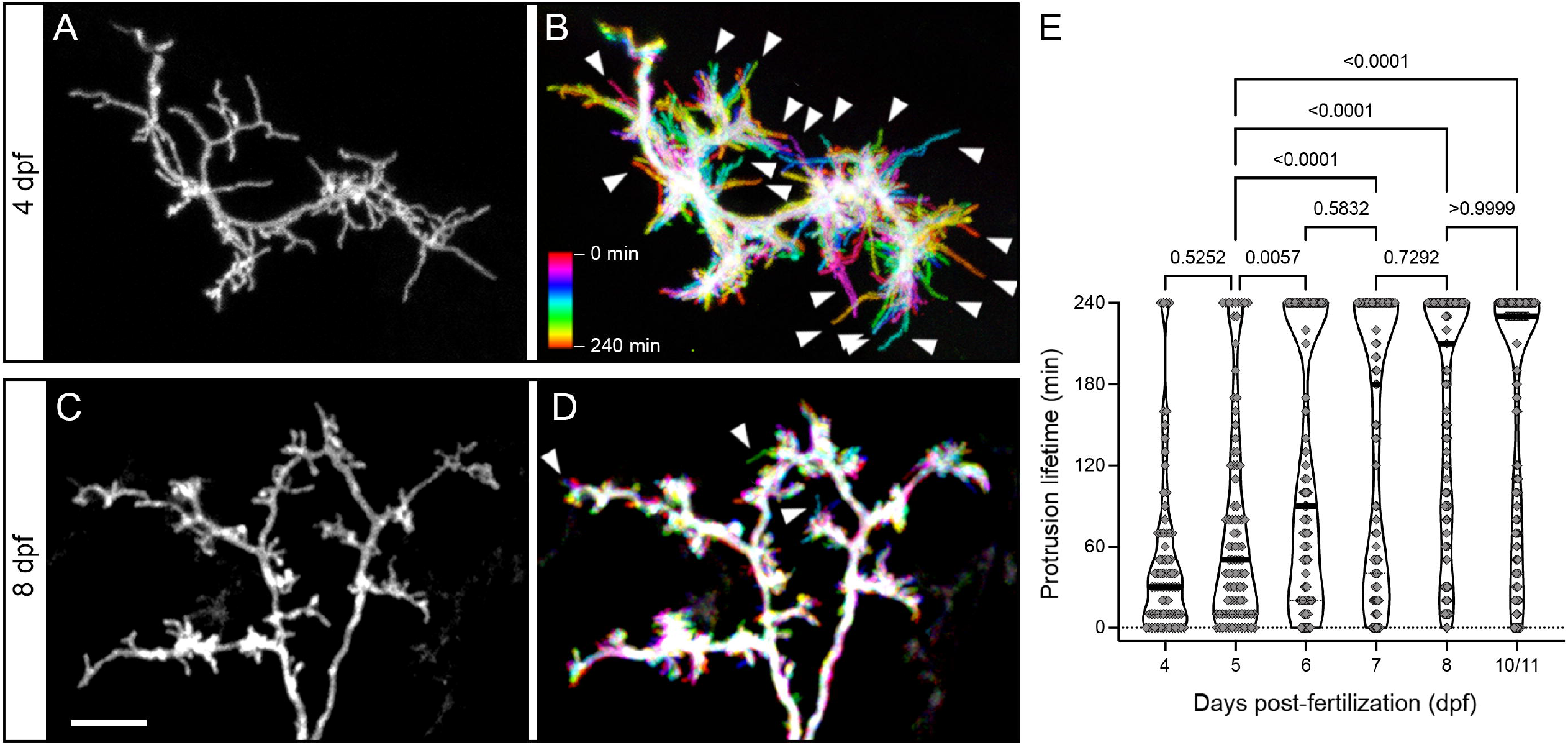
Developmental changes in protrusion stability. **A**. High magnification view of the SM-targeted apical dendrite of a 4 dpf PyrN. Note abundance of long, filopodial protrusions. **B**. Temporal color coded image of dendrite in A during a 4 hr timelapse recording with image stacks acquired every 10 min. Color scale indicates which image corresponds to each timepoint. Note prevalence of motile protrusions with a single color label, indicating they were present at that position during a single acquisition (arrowheads). **C**. High magnification view of the SM-targeted apical dendrite of an 8 dpf PyrN. Note the abundance of short, spine-like protrusions. **D**. Temporal color coded image of dendrite in C during a 4 hr timelapse recording with image volumes acquired every 10 min. Note the relatively small number of filopodia-like motile protrusions generated (arrowheads). **E**. Comparison of protrusion lifetimes at 4, 5, 6, 7, 8, and 10/11 dpf. Note gradual increase in stable protrusions (240 min lifetimes) and decrease in transient spines lasting less than 120 min. Horizontal line on violin plot indicates median value for each group. One-way ANOVA with Tukey’s multiple comparisons posthoc test. Significant differences not shown on graph: 4 dpf vs. 6dpf: p<0.0001, 4 dpf vs. 7dpf: p<0.0001, 4 dpf vs. 8 dpf: p<0.0001, 4 dpf vs. 10/11 dpf: p<0.0001, 6 dpf vs. 8dpf: p=0.015, and 6 dpf vs. 10/11dpf p=0.0154. Number of spines and neurons analyzed at each timepoint: 73 (4), 91 (5), 99 (4), 83(4), 123 (4), and 110 (4). Scale bar: 5μm in A-D.

### PSD95-EGFP accumulation is weakly correlated with spine stability

To directly examine the relationship between spine stability and formation of synaptic contacts, we imaged PyrNs expressing the postsynaptic marker PSD95-EGFP and DsRed as a cytosolic marker (Niell et al., 2004). As previously reported (Demarco et al., 2021), high levels of PSD95-EGFP expression resulted in neurons with abnormal morphologies. The narrow range of expression levels yielding detectable fluorescence and normal PyrN morphologies caused us to exclude the majority of labeled neurons. This low yield precluded detailed analysis at multiple days of development. Therefore we focused on 6 dpf, an intermediate timepoint when PyrN apical dendrites have a mix of both stable and transient protrusions (Figure 4E). Consistent with our previous findings (Demarco et al., 2021), PyrNs contained a high density of PSD95 puncta in their apical dendritic arbor (Figure 5A). Within the apical dendrite, the majority of puncta were located within spine heads (filled arrowheads in Figure B-D). Preliminary examination also revealed that protrusions with PSD95-positive heads tended to be stable (filled arrowheads in Figure 5B-D), whereas protrusions lacking puncta tended to retract (open arrowheads in Figure 5B-D). We also observed instances where nascent protrusions formed a PSD95-EGFP puncta and became stabilized (red arrowhead in Figure B-D), as well as instances where a spine was retracted but its PSD95 puncta persisted (cyan arrowhead in Figure B-D).

**Figure 5.**
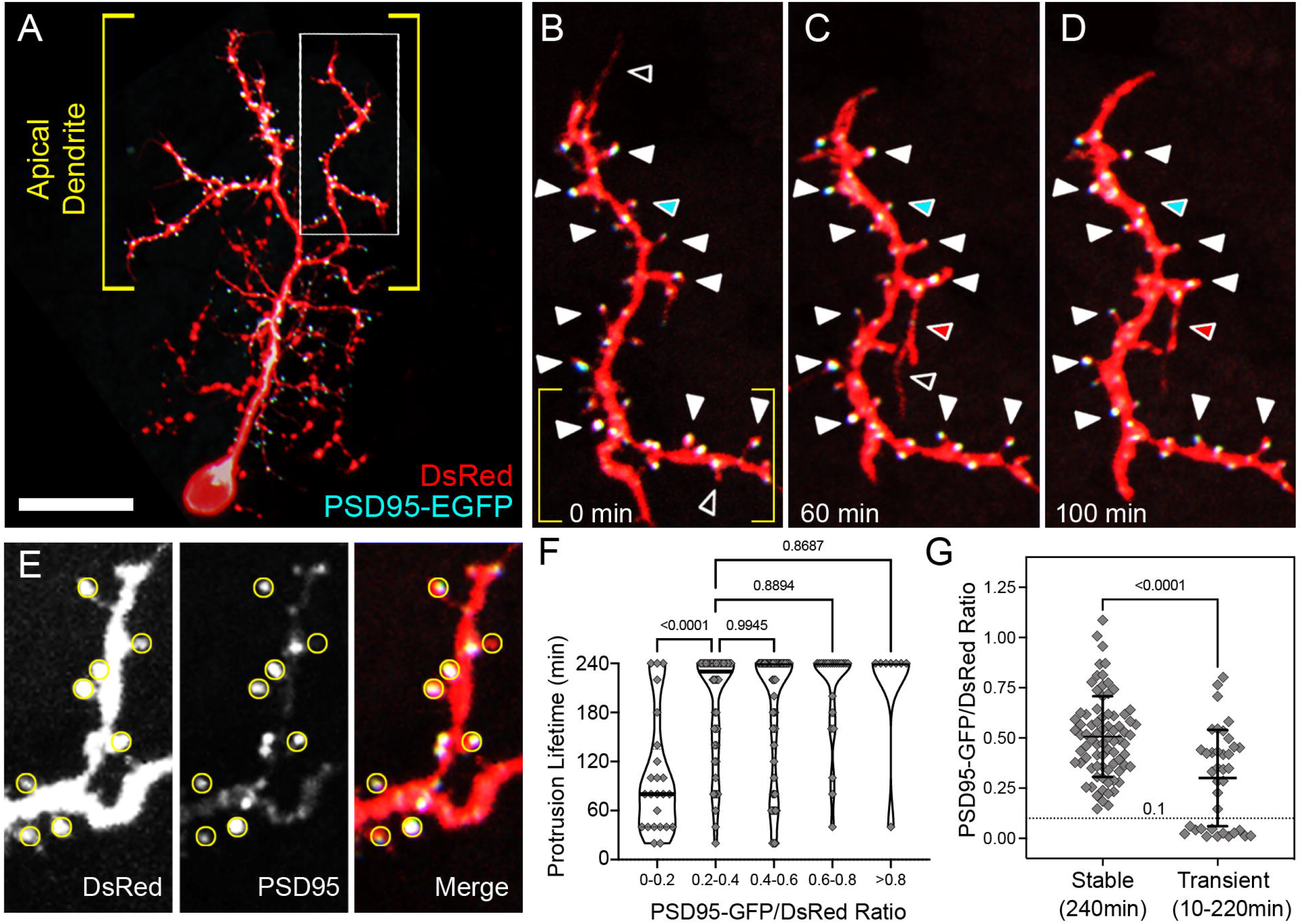
PSD95-EGFP dynamics in PyrN apical dendrites. **A**. Native orientation view of a 6 dpf PSD95-EGFP/DsRed labeled PyrN. **B-D**. Timelapse images of PSD95-EGFP localization in a dendrite subregion, as indicated by boxed region in A. White arrowheads mark seven PSD95-positive spines that were present during the entire 4 hr recording. For clarity some stable spines with PSD95 accumulations are not marked by arrowheads. Black arrowheads indicate PSD95-negative protrusions that were retracted during the timelapse. Red arrowheads indicate a protrusion that extended, formed a PSD95-EGFP puncta and was stabilized from 60 to 100 min. **E**. Magnified view of subregion indicated by yellow brackets in B. DsRed (left), PSD95-EGFP (middle), and merged (right) fluorescence channels are shown separately. Yellow circles indicate regions of interest used to calculate PSD95-EGFP/DsRed signal ratios. Note the wide range of PSD95-EGFP intensities within the analysis regions (middle and right panels). **F**. Quantification of protrusion lifetimes for spines binned into groups based on their degree of PSD95-EGFP enrichment (PSD95-EGFP/DsRed ratio). Note that there is a wide range of lifetimes for protrusions with the lowest PSD95-EGFP enrichment (ratios from 0-0.2), whereas the majority of protrusions with intermediate (0.2-0.6) or high (>0.6) PSD95-EGFP enrichment values were stable during the 4 hr recording. One-way ANOVA with Tukey’s multiple comparisons posthoc test. Significant differences not shown on graph: 0-0.2 vs 0.4-0.6: p<0.0001, 0-0.2 vs 0.6-0.8: p<0.0001, 0-0.2 vs >0.8: p<0.0001. Analysis performed on 156 protrusions from 5 neurons. **G**. Comparison of PSD95-EGFP/DsRed ratios between stable and transient protrusions. Note that both groups have many spines with intermediate values, but only the stable group has several spines with ratios > 0.8. Conversely, only the transient group contains protrusions with ratios <0.1. Unpaired t-test. Scale bar: 20μm in A, 8μm in B-D, 5μm in E.

To examine the relationship between PSD95 enrichment and protrusion stability, we measured the fluorescence intensity ratio between PSD95-GFP and DsRed at the tip of each protrusion at the first timepoint of each 4hr time-lapse recording (Figure 5E). Subsequently, we measured the lifetime of each protrusion to determine if there was a correlation between PSD95 enrichment and stability. This analysis revealed a statistically significant difference between the stability of protrusions with a low PSD95:DsRed ratio (0-0.2) versus those with higher ratios (p≤0.0006 for all pairwise comparisons, one-way ANOVA with Tukey’s multiple comparisons test; Figure 5F). However, there was not a linear relationship between the PSD95:DsRed ratio and protrusion lifetime. For example, protrusions with intermediate PSD95:DsRed ratios (0.2-0.4,) had similar average lifetimes as those with higher ratios (0.4-0.6, 0.6-0.8, and ≥0.8; p values >0.87; Figure 5F). To further examine this effect, we grouped protrusions by their lifetime (stable or transient) and plotted their corresponding PSD95:DsRed ratios (Figure 5G). This analysis revealed a highly significant difference in the average PSD95:DsRed ratio for transient vs. stable protrusions (p<0.0001, unpaired t test), in part due to the transient nature of every protrusion with a very low PSD95:DsRed ratio (≤0.1; Figure 5G). 96.2% (75 of 78) of stable spines with lifetimes of 240 min contained PSD95:DsRed ratios >0.2, although a considerable fraction of transient spines did as well (64.9%, 24 of 37). These findings support a model in which a minimum amount of PSD95 is required for stabilization (≥0.2 in our experiments). However, even relatively high levels of PSD95 enrichment do not necessarily protect spines from retraction. This is likely due to PSD95 being an early marker of postsynaptic contacts. Long-term spine stabilization likely requires recruitment of additional PSD components, such as glutamate receptors and cytoskeletal proteins.

### Impaired apical dendrite growth in *fmr1* mutant larvae

To examine dendrite development in *fmr1* mutant larvae, we generated *Tg(id2b:gal4)* fish harboring the *hu2787* mutation of *fmr1* (see methods; Broeder et al., 2009). Embryos generated from incrosses of these fish were injected with *uas:egfp-caax* plasmid DNA to label isolated PyrNs. Live imaging of EGFP-caax labeled neurons in 8dpf *fmr1* mutant larvae revealed normal PyrN dendrite stratifications in SM, SFGS, and SGC layers of tectum (Figure 6A-B). These image volumes also consistently revealed smaller and less complex apical dendrites in *fmr1* mutant larvae (Figure 6A-B). To quantify this effect, we used the SNT (Simple Neurite Tracer; Arshadi et al., 2021) plugin for FIJI/ImageJ to semi-automatically generate skeletonized 3D tracings of PyrNs in WT and *fmr1* mutants. These tracings were used to measure neurite length in 3D for SM-, SFGS-, and SGC-stratified PyrN arbors. Quantification of neurite length measurements revealed significant decreases in both SM and SGC arbors in *fmr1-/-*larvae (Figure 6E). To quantify the synaptic territory of apical dendrites, we calculated the area of each arbor along the retinotopic axes using maximum projections of image subvolumes containing a single PyrN arbor (Figure 6C-D; also see Figure 1B). Retinotopic area measurements revealed a significant decrease only in the SM apical PyrN dendrite in *fmr1-/-*mutants. Decreased SGC arbor size without a decrease in retinotopic area suggests the primary effect of *fmr1* loss on this arbor is reduced branching. The SGC arbor contains both axonal and dendritic segments (DeMarco et al., 2019; Demarco et al., 2021). Loss of *fmr1* reduces axon arbor complexity in mouse barrel cortex (Bureau et al., 2008), therefore this effect may be due to disruption of SGC axon branching. The reductions in neurite length and retinotopic area of SM dendrites indicate that loss of FMRP impairs both dendritic growth and branching of the spiny PyrN apical dendrite. The lack of effect on the SFGS dendrite, which does not form spines, suggests *fmr1* may play a more critical role in the growth and synaptogenesis of spiny dendrites with a high input density.

**Figure 6.**
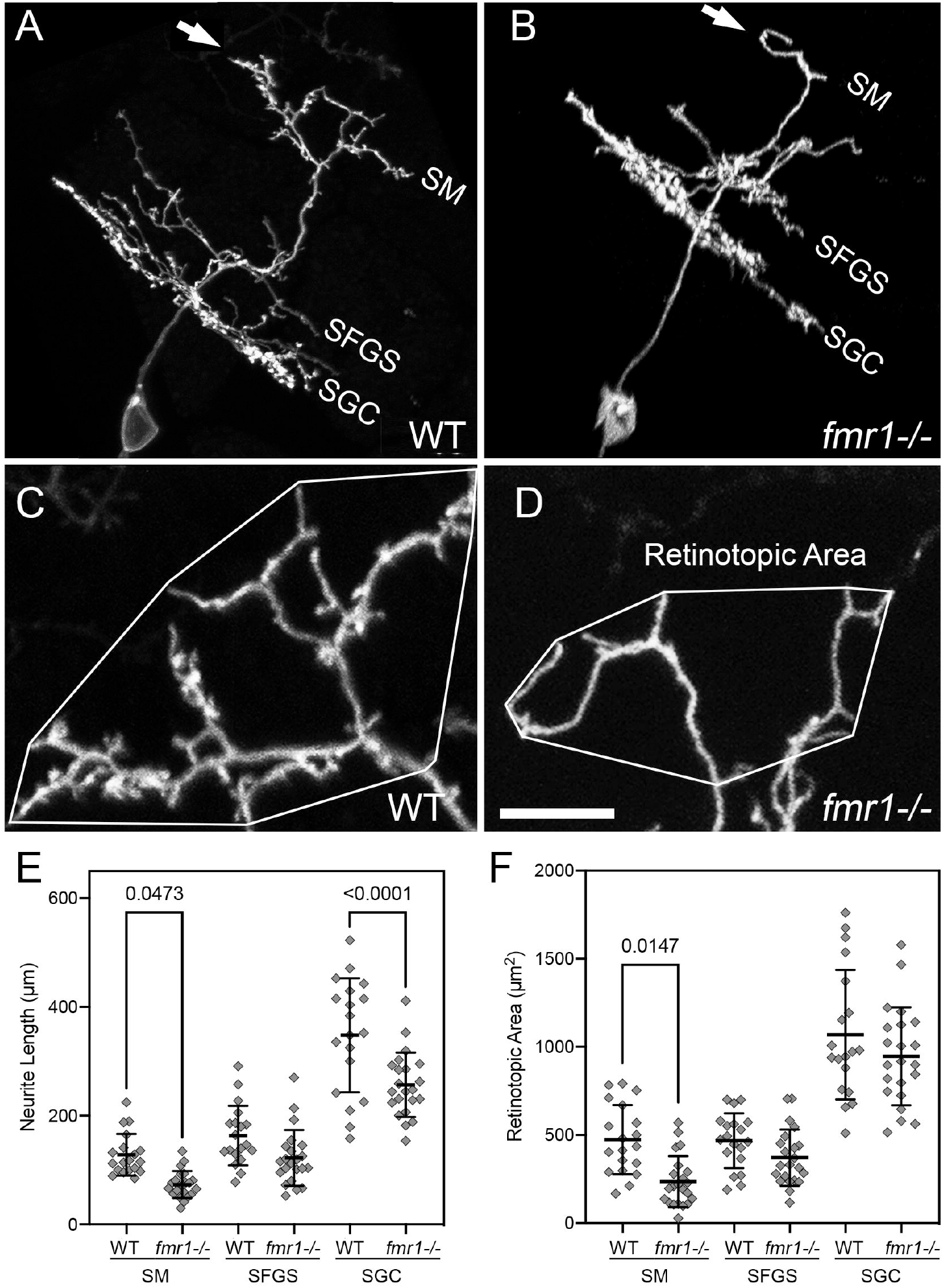
*fmr1* mutants exhibit defects in PyrN dendrite development. **A-B**. Sideview rotated image volumes of a PyrN in a WT 8dpf larva (A) and an 8dpf *fmr1*-/-larva. Note three distinct dendrite stratifications in layers SM, SFGS, SGC. Arrows in A and B indicate the apical dendrite. **C-D**. Native orientation view (dorsal side up as in Figure 1B) of the SM-targeted apical dendrites in the two PyrNs shown in A-B. Convex polygons overlayed on the arbor demonstrate how retinotopic area was calculated. Note the reduction in dendrite length and area, as well as very few spines on the *fmr1* mutant dendrite. **E**. Dendrite arbor-specific neurite length measurements in WT vs. *fmr1* mutants. Note significant reductions for the SM and SGC arbor in the mutant. One-way ANOVA with Tukey’s multiple comparisons posthoc test. **F**. Retinotopic measurements in WT vs. *fmr1* mutants. Note significant reduction only for the SM dendrite arbor in *fmr1* mutants. One-way ANOVA with Tukey’s multiple comparisons posthoc test. Analysis performed on 19 WT and 22 fmr1 mutant PyrNs. Scale bar, 15μm in A-B, 5μm in C-D.

### Loss of *fmr1* leads to reduced spine densities and immature spine morphologies

Spine density and morphology were examined in EGFP-caax labeled PyrNs at 8dpf, a timepoint when the PyrN apical dendrite contains mostly short spines with enlarged heads (Figure 3). Structural imaging of WT dendrites revealed a dense array of spines with mature morphologies (Figure 7A). In contrast, *fmr1* mutant dendrites had a sparse distribution of long, filopodia-like protrusions with narrow heads (Figure 7B-C; see also Figure 6d). This trend was observed in mutant dendrites with severely impaired dendrites (Figure 7C) as well as those with moderate reductions in dendritic arbor size (Figure 7B). On average, PyrNs in *fmr1* mutants exhibited a greater than 3-fold reduction in spine density (Figure 7D; 1.573±0.46 vs. 0.477±0.16 μm^-1^, n=17 neurons for each condition, p value <0.0001, unpaired students t-test). This suggests PyrNs in *fmr1* mutants are impaired in their ability to form or stabilize spines, as either could lead to a reduction in density. Spine head width was also significantly reduced in PyrNs of *fmr1* mutant larvae (Figure 7E), indicative of more spines with immature morphologies. Developmentally, there is a correlation between spine head width and spine stability (Figures 3 and 4), suggesting reduced spine density in *fmr1* mutants is caused by a specific deficit in spine stabilization.

**Figure 7.**
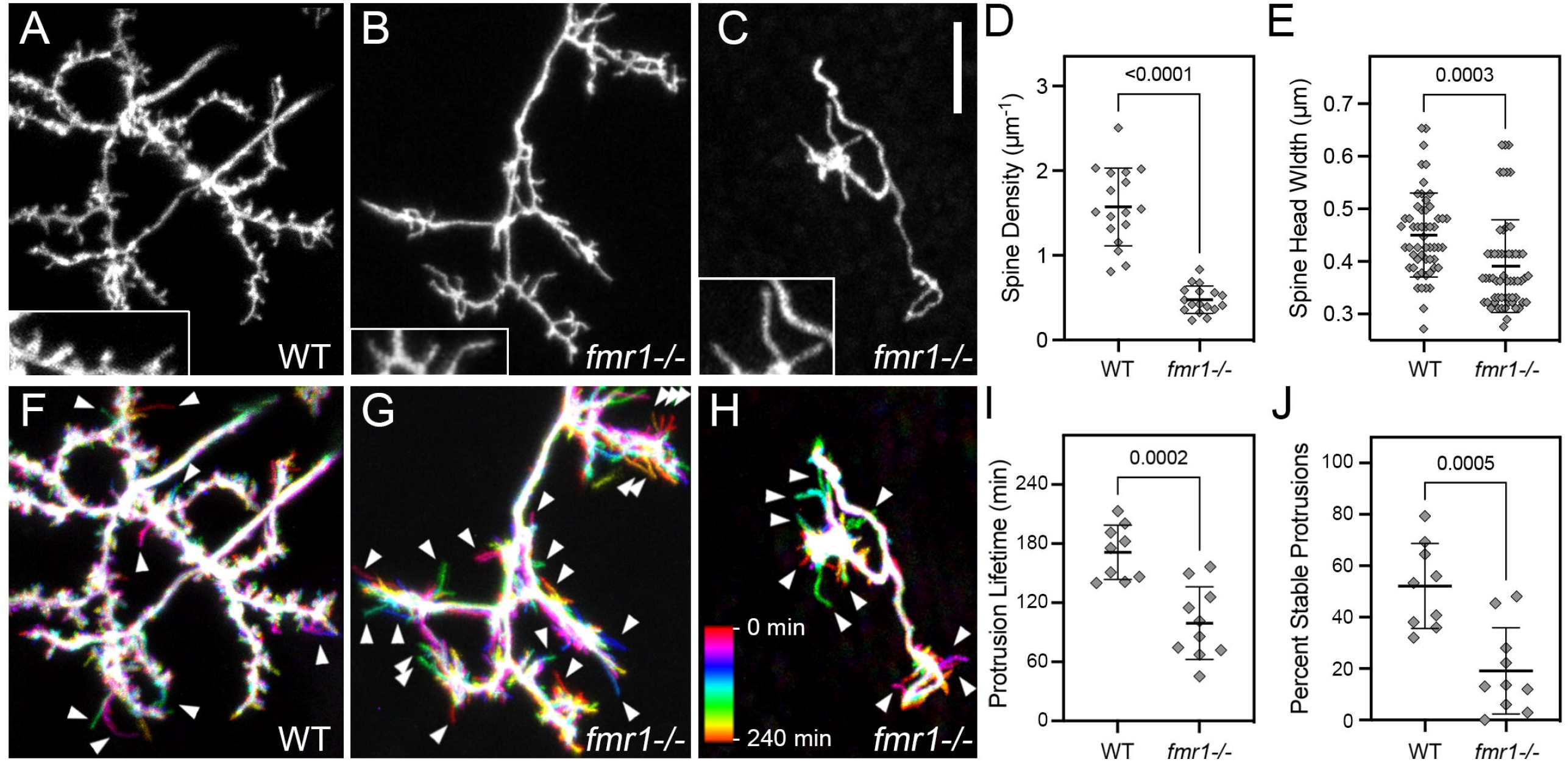
Reduced spine density, head width, and stability in *fmr1* mutants. **A**. Native orientation maximum projection images (dorsal side up as in Figure 1B) of the SM-targeted apical dendrites in a 8dpf WT larva. **B**. Native orientation views of the SM-targeted apical dendrites from two 8dpf *fmr1* mutant larvae. These are representative of the mild (B) and severe (C) phenotypes observed in the *fmr1* mutants. Insets in A-C are 1.5x magnified views of subregions containing spines. Note that the WT has both thin and wide heads, whereas the majority of spines on *fmr1*-/-PyrNs have thin heads. **D-E**. Quantification of spine density and spine head width. Both measurements were made using static single maximum projection images as those shown in (A-C). Note significant decreases in both density and head width in the *fmr1* mutants. **F-H**. Temporal color coded image of dendrites in A-C during 4 hr timelapse recordings with image volumes acquired every 10 min. Color scale in H applies to all three images and indicates which image corresponds to each timepoint. Note increased number of transient protrusions (single color label indicating they were present during a single timepoint) in *fmr1* mutants compared to WT (arrowheads). **I-J**. Quantification of protrusion lifetime and percent stable protrusions. Both measurements were made using 4hr timelapse recordings with a 10 min acquisition interval. On average PyrN apical dendrites in *fmr1* mutant larvae had reduced protrusion lifetimes (I), an effect largely due to a significant reduction in the percentage of stable protrusions present throughout the entire 4hr timelapse. In D, E, I, and J all p values were obtained using unpaired t-tests. Scale bar, 5μm.

### Spine stabilization is impaired in *fmr1* mutant larvae

To determine if the morphological changes observed in PyrN spines of *fmr1* mutants correlate with changes in spine dynamics, we performed time-lapse imaging of EGFP-caax labeled PyrNs at 8dpf. Inspection of these recordings revealed dendrites in *fmr1* mutants with increased turnover of spines compared to WT (Supplemental Movie 2). To capture this difference in static images, we generated color-coded temporal projections from these recordings (Figure 7F-H). WT dendrites formed few motile, filopodia-like protrusions (arrowheads in Figure 7F), whereas in *fmr1* mutants protrusions were predominantly motile (arrowheads in Figure 7G-H). Lifetime analysis confirmed a significant reduction in average protrusion lifetime in *fmr1* mutants compared to WT (Figure 7I). This was in large part due to a marked reduction in the percentage of protrusions with lifetimes of 240 min in *fmr1* mutants (19.3±16.6% in *fmr1* mutants vs. 52.3±16.8% in WT, n=10 and 9 dendrites, p=0.0005, unpaired t-test). These data support a model in which immature spine morphologies in *fmr1* mutants reflect an inability to form stable synaptic contacts.

## DISCUSSION

Our findings confirm many similarities between spine development in zebrafish and mammals. Morphologically, spines in mature PyrNs (older than 6dpf) exhibited a wide range of morphologies, from filopodia-like structures with thin heads to short spines with mushroom heads (Figure 1). The filopodia-spine transition in PyrNs was characterized by a gradual shift in morphologies, from long filopodia at 4-5 dpf to short spines with enlarged heads at 8-11dpf (Figure 3). A similar progression occurs in mammalian pyramidal cell dendrites (Dailey and Smith, 1996; Maletic-Savatic et al., 1999; Ziv and Smith, 1996), although this process occurs more rapidly in zebrafish PyrNs. A unique feature of this system is the ability to examine dendrite branch remodeling in relation to spine development. In mammalian pyramidal cells, these processes are generally not concurrent. In organotypic slices of rat hippocampus, pyramidal cell dendrites undergo an early developmental stage (1-2 d in culture) where transient filopodia predominate and can transform into new dendritic branches (Dailey and Smith, 1996). However, at timepoints when short, stable spines predominate (1-2 weeks in culture), dendritic arbors do not exhibit changes in branch number or length (Dailey and Smith, 1996). Between 6 and 11dpf, PyrN dendrites are forming spines yet continue to undergo large-scale changes in dendrite morphology (growth, branching, and retraction; see Figure 2). The rapid development of the zebrafish visual system (Niell and Smith, 2005) may drive an accelerated rate of synaptogenesis relative to dendrite development in PyrNs. Alternatively, the dendritic rearrangements during this period may reflect retinotopic refinements necessitated by the continued growth of both retina and brain. We have previously observed changes in the retinotopic position of RGC axons arbors in the tectum during this period of development (Robles et al., 2013). As both retina and brain continue to grow throughout the life of zebrafish, such rearrangements could be expected to persist into adulthood.

PyrN apical dendrites form a dense array of glutamatergic postsynaptic specializations containing PSD-95 (Demarco et al., 2021). Time-lapse analysis of PSD95-EGFP dynamics allowed us to define the relationship between spine lifetime and PSD95-EGFP accumulation. Our data indicate that the vast majority (96.2%) of stable spines with lifetimes ≥4hrs contain discrete PSD95 accumulations. Although spines lacking PSD95-EGFP accumulations were generally short-lived, there was only a weak correlation between PSD95-EGFP enrichment and spine stability. This is evidenced by the similar mean lifetimes of spines with high or intermediate PSD95-EGFP/DsRed ratios (Figure 5). These results are similar to previous findings in mouse cortical pyramidal neurons, where PSD95 accumulations can be present in both short-lived and stable spines (Cane et al., 2014). Together, these findings support a model in which PSD95 is an early marker of synapses formed by spines, labeling both short-lived and long-lived spines. This early role in synapse formation is likely to involve PSD95’s well-characterized role in recruiting and clustering AMPA and NMDA-type glutamate receptors (Chen et al., 2015).

Our finding that *fmr1* mutants exhibit reductions in spine stability is consistent with previous observations in mouse cortex (Cruz-Martín et al., 2010; Pan et al., 2010). In both of these studies FMRP-deficient dendrites contained a higher percentage of transient protrusions compared to WT controls. In mouse layer 5 pyramidal neurons, transient spines were also shown to have reduced spine head widths and volumes (Pan et al., 2010). Although our data revealed a reduction in spine head width for PyrNs in *fmr1* mutant larvae, this effect on spine head width may be cell type-specific. For example, in FMRP-deficient purkinje cells, spine immaturity is reflected in increased spine lengths without a change in head width (Koekkoek et al., 2005). Unlike our study and others in *Fmr1* KO mice (Comery et al., 1997; Nimchinsky et al., 2001), Purkinje cells also did not exhibit a change in spine density or dendritic arbor complexity. One possible explanation is that FMRP plays a more critical role in circuits where spiny neurons undergo high levels of activity-dependent synapse refinement. Purkinje dendrite arborization and synaptogenesis may be genetically specified to a greater extent than cortical neurons and tectal PyrNs. If severity of phenotype in *fmr1* mutants reflects dependence on activity-dependent competition, this would suggest tight control of PyrN dendrite arborization by activity-dependent mechanisms.

Although we have characterized the time-course of spine development in PyrN apical dendrites, we do not know if this process is activity dependent. PyrNs are visually responsive neurons comprised of three functional classes encoding different luminance ranges during luminance fluctuations: ON, OFF, and DUAL (Tesmer et al., 2022). Our current model of PyrN activation predicts TL inputs to PyrN apical dendrites to be class-specific (ON, OFF, DUAL), matching the response class of RGC input to the medial dendrite. But how do PyrN apical dendrites selectively wire up with TL inputs with matching responses? Establishment of the tectal neuropil layers is largely independent of neural activity (Nevin et al., 2008) and largely controlled by layer-specific guidance cues (Xiao and Baier, 2007; Xiao et al., 2011). These guidance cues likely guide PyrN apical dendrites to the tectal surface in an activity-independent manner, as evidenced by the rapid formation of layer-specific dendritic stratifications (Figure 2). Apical dendrite stratification is followed by a period of dendritic growth, branching, and spine formation starting between 3 and 4 dpf (Figures 2, 4, and 5). The earliest visual responses in tectum begin at 2.75 dpf (Niell and Smith, 2005), soon after RGC axons first innervate tectum. If RGC input to PyrNs is established during this early phase, spine synapses on PyrN apical dendrites could form by an activity-dependent mechanism in which synaptic activity at each spine is compared to spike patterns driven by RGC input. Excitatory postsynaptic potentials (EPSPs) triggered by matched inputs (ON, OFF, or DUAL) would coincide with backpropagating action potentials capable of relieving the Mg2+ block of NMDARs, thereby boosting Ca^2+^-influx in a spine-specific manner. NMDAR-dependent Ca2+ influx is known to play a critical role in spine stabilization and enlargement in many circuits (Nikonenko et al., 2002). Conversely, mismatched inputs with small synaptic Ca2+ influx events would be weakened and disassembled. Future experiments will be aimed at determining if activity blockade in PyrNs disrupts PyrN apical dendrite spine formation.

The strong defects we observe in dendrite development and spine stabilization may seem surprising considering the mild behavioral defects described in the *hu2787 fmr1* mutant (Ng et al., 2013). One possibility is that only certain types of neurons have a stringent requirement for normal levels of FMRP during development, which makes them more susceptible to its loss. Tectal PyrNs form part of a reciprocal circuit between the tectum and TL. The apical dendrite in SM layer of tectum receives excitatory input from TL (Folgueira et al., 2020; Northmore, 2017), and our recent anatomical data suggests a high degree of input convergence at the TL-PyrN synapse (Demarco et al., 2021). TL axons providing input to the PyrN apical dendrite in SM form extremely large axonal arbors with a high degree of overlap. These large synaptic territories distribute information from each TL neuron to PyrNs located throughout tectum. In stark contrast to TL axons, PyrN apical dendrites are small and contain as many as 100 postsynaptic specializations (Demarco et al., 2021). High input densities onto PyrN apical dendrites may necessitate spines, which increase dendritic surface area and biochemically isolate synapses. We propose that spiny dendrites with high input densities are more susceptible to loss of FMRP than aspiny dendrites. Consistent with this possibility, loss of FMRP had no effect on neurite length or area of the medial (SFGS) PyrN dendrite that receives input from RGC axons.

Could spiny dendrites that rapidly form many spines be particularly sensitive to loss of FMRP? Presynaptic TL axons form a dense, mesh-like plexus in the SM layer of tectum (Demarco et al., 2021). One possible explanation for PyrN sensitivity to loss of FMRP is that a high density of presynaptic terminals requires dendrites to simultaneously sample many different potential partners. Consistent with this, protrusion density remains relatively high throughout the developmental period we examined (Figure 2K). This may create an increased need for early components of PSDs, such as scaffolding proteins and glutamate receptors. In mouse cortex and hippocampus, FMRP binds to mRNAs encoding several components of the PSD, including scaffolding proteins of the Shank, MAGUK, and SAPAP families, as well as subunits of NMDA-type and AMPA-type glutamate receptors (Schütt et al., 2009). At the protein level, loss of FMRP reduced levels of the PSD protein SAPAP3, while increasing levels of SAPAP1, glutamate ionotropic receptor AMPA type subunit 1 (GRIA1), and glutamate ionotropic receptor NMDA type subunit 1 (GRIN1). However, this study also found cell type-specific effects on protein levels when comparing cortical and hippocampal neurons. Disruption in the balanced levels of synaptic proteins might disrupt normal synapse assembly and/or disassembly. We propose the effect on apical dendrite arborization to be secondary to spine stabilization deficits in *fmr1* mutants. Synaptotropic models of dendrite growth propose stable synaptic contact biases the direction of growth by protecting a subset of branches from retraction while others are retracted (Cline and Haas, 2008; Niell, 2006). From this view, the mature dendritic arbor morphology reflects the location of strong connections with appropriate presynaptic partners. Thus, impaired dendrite growth in *fmr1* mutant PyrNs likely arises due to the neuron’s inability to form strong synaptic contacts and thus protect nascent branches from.

In conclusion, *in vivo* imaging of dendritic spine morphogenesis in zebrafish larvae was enabled by examining a genetically defined neuron type in optic tectum. In addition to confirming similarities between spine morphogenesis in zebrafish larvae and mammals, these findings establish the larval zebrafish as a valuable animal model to study cellular mechanisms underlying spine defects in neurodevelopmental disorders. Although in this study we have focused on a zebrafish model of Fragile X, there are currently more than a dozen validated zebrafish lines with mutations in paralogs of ASD risk genes (Rea and Van Raay, 2020). The ability to conduct automated, high-resolution brain imaging in living zebrafish larvae (Early et al., 2018; Pardo-Martin et al., 2010) will enable future studies to monitor PyrN spine morphology and dynamics with increased throughput. Several scalable screening approaches possible using this approach include: high throughput drug discovery (MacRae and Peterson, 2015), forward genetic screening (Muto et al., 2005), and reverse genetic screening using CRISPR (Shah et al., 2015) techniques.

## Materials and Methods

### Fish Lines

Zebrafish adults and larvae were maintained at 28°C on a 14/10 h light/dark cycle. *Tg(id2b:Gal4-VP16)mpn215* and *Tg(UAS-E1B:NTR-mCherry)c264* transgenic lines have been previously described (Davison et al., 2007; Förster et al., 2017). All larvae used were either mutants for *mitfa*^*-/-*^ *(nacre) or* double mutants for *mitfa*^*-/-*^ and *roy*^*-/-*^ (*casper*). Use of these pigmentation mutants eliminated the need to chemically block skin pigmentation with phenythiourea (PTU). All animal procedures conformed to the institutional guidelines of the Purdue University Institutional Animal Care and Use Committee (IACUC). *fmr1*^*hu2787*^ mutant fish (Broeder et al., 2009, 1) were obtained from the Zebrafish International Resource Center (ZIRC).

### Genotyping

Adult and larval fish were genotyped as described previously (Ng et al., 2013). Briefly, we used the following PCR primers: forward primer (5′-CTA AAT GAA ATC GTC ACA TTA GAG AGG GTA) and reverse primer (5′-TCCATG ACA TCC TGC ATT AG). PCR products were digested with RsaI restriction enzyme to identify WT and homozygotes. *fmr1*^*hu2787*^ mutant fish in the A/B strain were mated with *Tg(id2b:gal4,uas:NTRmcherry)* fish in the *mitfa*^*-/-*^/*roy*^*-/-*^ background. Heterozygotes were identified in the subsequent generation and mated with WT *casper* fish to generate heterozygotes carrying the *id2b:gal4* transgene. These heterozygotes were in-crossed for embryo injections and larvae with sparse labeling were imaged and then genotyped by PCR. Uninjected embryos from these incrosses were also reared to generate adult homozygotes, which were healthy and exhibited normal fecundity. Approximately half of *fmr-/-*mutant data in this study were obtained from in-crosses of *fmr-/-*adult fish.

### Embryo Injections

Genetic mosaic labeling of single neurons by expression of a membrane targeted EGFP was achieved by injection of the 4xnrUAS:EGFP-caax plasmid (a gift from B. Appel and J. Hines, University of Colorado, Denver, CO), along with RNA encoding Tol2 transposase into WT or mutant embryos transgenic for *id2b:Gal4-VP16*. Labeling of single neurons by co-expression of a PSD95-EGFP fusion protein and DsRed as a cytosolic marker was achieved by injection of the 14UAS PSD95:GFP 5UAS DSRedExpress plasmid (addgene plasmid #74315). All DNA constructs were pressure-injected at a concentration of 25–50 ng/l into one-to eight-cell-stage embryos.

### Confocal Imaging

For live confocal imaging between 4 and 11 dpf larvae were anesthetized in 0.016% tricaine and embedded in 2% low-melting-point agarose. Imaging was performed on a Nikon C2 confocal microscope equipped with solid state lasers for excitation of EGFP (488 nm) and mCherry/TagRFP (555 nm). Whole-brain imaging of larvae was performed using a Nikon LWD 16x 0.8NA water immersion objective using 1-1.5 μm z-steps. Larvae with single labeled neurons were imaged using a Nikon 60x 1.0NA water immersion objective and 0.375-0.5 μm z-steps. For timelapse recordings, laser power was lowered to <1% and z-stacks with 0.6-1μm z-steps were acquired every 10 min for 4 hrs, yielding 25 image volumes. For PSD95-EGFP/DsRed imaging, z-stacks were acquired every 20 min for 4 hrs to prevent photodamage due to the increased number of scans per z-stack due to two-channel acquisition. Additionally, prior to each timelapse acquisition, laser power and detector gain for both channels were adjusted to yield images in which spines with bright PSD95-EGFP puncta had PSD95-EGFP/DsRed intensity ratios close to 1.0. This circumvented the need to normalize ratio values when combining data from multiple neurons (Figure 4F-G).

### Image processing and analysis

Image stacks were visualized and analyzed using ImageJ FIJI software (http://fiji.sc/Fiji). 3D rendering was performed using the 3D Viewer plugin (Schmid et al., 2010). Skeletonized tracings used for calculating neurite lengths were generated with the semi-automated neurite segmentation plugin SNT/Simple Neurite Tracer (Longair et al., 2011). Arbor specific neurite lengths could be automatically measured in 3D. Retinotopic area was defined as the convex hull area for each arbor when viewed in the native orientation as in Figure 1A-B (Teeter and Stevens, 2011). Protrusion density was calculated by manually annotating spine locations on maximum projections of apical dendrite image volumes. Protrusion lengths and lifetimes were measured manually in ImageJ/FIJI. Protrusion head widths were measured by obtaining intensity profiles from regions spanning the distal 0.5 μm of each spine as shown in Figure 3F, H, and J. To ensure that bends or swellings of the dendrite were not counted as protrusions, only protrusions with lengths >0.5 μm were analyzed for head width, density, or lifetime. Based on our length analysis presented in Figure 3I, 7.5% of protrusions (79 of 1,043) had lengths >0.5 μm. This is the potential undercount due to our length criteria, although it should be noted that this criteria was applied uniformly to control and experimental datasets. PSD95-EGFP/DsRed intensity ratios were measured on the first maximum projection of each timelapse sequence using a circular ROI with a diameter of 0.5μm. All data are presented as mean ± SD, except for violin plots where the horizontal line indicates the median value.

### Statistical Analysis

Data sets were analyzed using GraphPad Prism software version 9.2.0 for Mac (GraphPad Software, Inc., La Jolla, CA). All data displayed a normal distribution. One-way ANOVA was used to identify differences among means for data sets with three or more groups, with Tukey’s multiple comparisons test used posthoc for comparisons between groups. P-values less than 0.05 were considered significant.

## Acknowledgements

The authors thank Alejandra Agredo and Emily Malek for initial protocol testing and optimization.

## Data Availability

All data supporting the main conclusions of this paper will be included in this published article as supplementary information files. Additionally, data files and neuron tracings will be made available through the Purdue University Research Repository (purr.purdue.edu) in a dataset associated with the DOI: (pending).

## Figure Legends

**Supplemental Movie 1. The filopodia-spine transition revealed by live imaging of dendrite protrusion dynamics**. Timelapse recordings of the same EGFP-caax labeled PyrN apical dendrite imaged at 4, 6, and 8 dpf. Image volumes were acquired every 10 min for 4 hrs. Note prevalence of long, motile filopodia at 4 dpf, whereas short spines predominate at 8 dpf. At 6 dpf the dendrite contains a mix of long and short protrusions.

**Supplemental Movie 2. Reduced protrusion stability in *fmr1* mutant PyrNs**. Timelapse recordings of the three EGFP-caax labeled PyrN apical dendrites from one WT (left) and two *fmr1* mutant (middle and right) larvae. *fmr1* mutant PyrN at middle exhibited a moderate reduction in dendrite arbor size, whereas *fmr1* mutant PyrN at right exhibited severely impaired dendrite growth. Compare short and stable spines on WT to the motile filopodia present on the mutant PyrNs. Asterisk indicates a 1 hr interval when part of the middle dendrite was below the image acquisition volume due to specimen drift.

